# Contactless recordings of retinal activity using optically pumped magnetometers

**DOI:** 10.1101/2021.05.18.444672

**Authors:** Britta U. Westner, James I. Lubell, Mads Jensen, Sigbjørn Hokland, Sarang S. Dalal

## Abstract

Optically pumped magnetometers (OPMs) have been adopted for the recording of brain activity. Without the need to be cooled to cryogenic temperatures, an array of these sensors can be placed more flexibly, which allows for the recording of neuronal structures other than neocortex. Here we use eight OPM sensors to record human retinal activity following flash stimulation. We compare this magnetoretinographic (MRG) activity to the simultaneously recorded electroretinogram of the eight participants. The MRG shows the familiar flash-evoked potentials (a-wave and b-wave) and shares a highly significant amount of information with the electroretinogram recording (both in a simultaneous and separate recording). We conclude that OPM sensors have the potential to become a contactless alternative to fiber electrodes for the recording of retinal activity. Such a contactless solution can benefit both clinical and neuroscientific settings.

## 1 Introduction

Optically pumped magnetometers (OPMs) have attracted increasing attention in neuroscientific research over the last few years (e.g., Alem et al., 2014; Boto et al., 2017, 2018; Hill et al., 2019; Iivanainen et al., 2020; Roberts et al., 2019; Zetter et al., 2018). Without the need to be cooled to cryogenic temperatures like superconducting quantum interference device (SQUID) magnetometers, OPMs require relatively minimal insulation and do not need to be mounted at fixed locations within a dewar like traditional MEG. The sensors can rather be placed individually and close to the scalp, which has yielded better signal-to-noise ratio (SNR) for recordings of brain activity in simulations (Boto et al., 2016; Iivanainen et al., 2017) and real recordings (Boto et al., 2017, 2018; Hill et al., 2020; Iivanainen et al., 2020). The individual placement possibilities of the OPM sensors has also inspired novel placements, e.g. in the mouth for recordings of hippocampal activity (Tierney et al., 2021).

In this study, we exploit these new possibilities for the recording of retinal activity. Electrophysiological recordings of the retina are used in research (e.g., Bach et al., 2018; Heinrich and Bach, 2004, 2002) and clinical settings (Marmor et al., 1989, 2009; McCulloch et al., 2015) and are performed with either a contact lens electrode or a fiber electrode. The electroretinogram (ERG) reflects the summed activity of different cell populations in the retina and features slow evoked potentials as well as high-frequency oscillatory activity (Frishman, 2013). A light flash typically stimulates a negative potential around 25ms after flash onset, the *a-wave,* arising from photoreceptor activity (Perlman, 2001; Frishman, 2013). Following the a-wave, a positive wave around 80ms, the *b-wave,* is attributed to the activity of ON bipolar cells (Sieving et al., 1994; Vukmanic et al., 2014; Frishman, 2013). Concurrent with the a-wave is a high-frequency oscillatory burst (centered around 120 Hz), the *oscillatory potential* (Fröhlich, 1914; Munk and Neuenschwander, 2000). Its origin is attributed to diverse cell populations (Doty and Kimura, 1963; Kenyon et al., 2003; Frishman, 2013) and it is transmitted to visual cortex (Lopez and Sannita, 1997; Neuenschwander et al., 2002; Todorov et al., 2016; Westner and Dalal, 2019, but see Doty and Kimura, 1963; Heinrich and Bach, 2004; Molotchnikoff et al., 1975).

The recording of retinal activity is typically done with electrodes, however, the corresponding magnetic activity can be recorded instead, as pioneered by Aittoniemi et al. (1979) and Katila et al. (1981) using a SQUID sensor. I has also been reported that the frontal and temporal sensors of whole-head SQUID MEG systems can pick up retinal activity (Shigihara et al., 2016). The magnetoretinogram (MRG) does not require any direct contact of the magnetometer with the eye of the participant. OPMs give an exciting new opportunity to place several sensors around the eye of the participant, increasing both comfort compared to ERG recordings, and coverage compared to the early attempt at MRG recordings.

In this paper, we will show that OPM-MRG recordings of retinal activity are feasible and yield similar results to ERG recordings with fiber electrodes. This offers exciting new possibilities in vision research, visual neuroscience, and potentially also clinical applications.

## 2 Methods

### 2.1 Participants and recording setup

Nine participants took part in this study, however, the data from one participant had to be excluded due to an overly noisy electroretinogram (ERG) recording. Thus, we present the data of 8 participants (2 female; mean age 36.0 years, SD 3.97). All participants gave written informed consent and the study was approved by the Videnskabsetiske Komitéer for Region Midtjylland, Komitée II (Ethical Committees of Central Denmark Region, Committee II).

We used 8 OPM sensors (FieldLine Inc., Boulder, Colorado, USA; dimensions of one sensor: 1.3 × 1.5× 3 cm), which were spaced out as a 4 × 4 array in a 3D-printed holder (7.8× 9.4 cm). The measurements were done inside a 3-layer magnetically shielded room (Vacuumschmelze GmBh & Co. KG, Hanau, Germany). The participants were in a supine position and the OPM holder was mounted on a flexible arm to allow for a placement of the sensors close to the participants’ left eye. The holder did not have any physical contact with the participant in order to not introduce movement artifacts. The positioning of the holder lateral to the participant’s eye was guided by Katila et al. (1981) and the sensor grid was centered relative to the mid-line of the eye.

The ERG was recorded from the left eye of the participants using Dawson-Trick-Litzkow (DTL) fiber electrodes (Dawson et al. (1979); Diagnosys Vision Ltd., Dublin, Ireland). While the data from the OPM sensors was acquired with the FieldLine acquisition system, the ERG activity was recorded through the biopotential amplifier of the Elekta Triux MEG system (MEGIN Oy, Helsinki, Finland). A photo diode was mounted to the screen for stimulus onset definition and the data stream was recorded with both the OPM and MEG system for offline synchronization of the recordings.

To actively cancel the magnetic gradients produced by the magnetic parts of the Elekta MEG system situated in the shielded room, we used a Helmholtz coil around the magnetic parts of the MEG dewar. The gradients at the location of the measurement were nulled before the measurement started, using a fluxgate as feedback for manual adjustments of the currents sent to the Helmholtz coil. Immediately before the start of the recording, the OPM sensors measured the background fields in all three dimensions. Averaged across participants, the measured values had a range of 21.46 nT (—15.55 nT – 5.91 nT) in the *x* direction, a range of 1.97nT (—13.40nT —11.43nT) in the *y* direction, and finally a range of 4.72nT (—2.56nT – 2.17nT) in the *z* direction across all 8 sensors.

### 2.2 Stimuli

For visual stimulation, we presented brief flash stimuli to the participants. The flashes had a duration of 694 μs and a random inter-stimulus interval between 1000 and 1100 ms. The stimuli were projected onto a screen in front of the participant with a PROPixx projector (VPixx Technologies, Saint-Bruno, Canada) at 1440 Hz refresh rate. The stimuli had a brightness of 236 cd m^-2^ on the screen, and the room was dark during the presentation of the stimuli.

One set of stimuli consisted of 400 trials, and every participant viewed two sets. For the first set, the OPM sensors and the DTL fiber electrode were used for recording, for the second one, the fiber electrode was removed from the participant’s eye and only the OPM sensors were in place. This was done to have both simultaneously recorded data from ERG and MRG channels as well as a recording from the OPM sensors alone to ensure that any potential retinal activity measured by the OPMs was not introduced by the ERG electrode.

### 2.3 Data analysis

All data analysis was done in MATLAB using the open-source data analysis toolbox FieldTrip (Oostenveld et al., 2011) and custom scripts. All custom code used for data analysis is available at https://github.com/britta-wstnr/opm_mrg.

#### Basic processing

Epochs were created based on the photo diode signal to precisely identify flash onset. Line noise (50 Hz, 100 Hz, and 150 Hz) was removed using a Discrete Fourier Transform. Artifact rejection was done manually and for both data sets separately. For the simultaneous session with both ERG and MRG recorded, an average of 330.63 trials (SD 29.30) remained after artifact rejection. For the OPM-only data set an average of 380.38 trials (SD 19.73) remained.

The data were high-pass filtered at 1 Hz (onepass-zerophase, hamming-windowed FIR filter with order 1650) and low-pass filtered at 45 Hz (onepass-zerophase, hamming-windowed FIR filter with order 294). The data were then baseline corrected using the time window of —100 ms to 0 ms before flash-onset as baseline.

#### Mutual information

To quantify the shared information between the ERG and MRG, mutual information was computed within subjects between the ERG electrode and an OPM channel. The OPM channel was selected per data set and participant by searching for the highest b-wave amplitude across channels (search window for maximum: 0ms to 100 ms). The selection of one “best” channel per participant and data set was motivated by the fact that the placement of the OPM array could slightly vary per recording session and participant. Next, we computed the mutual information of the MRG channel with the ERG electrode for the time window of —50 ms to 150 ms per subject and data set. The computation of the mutual information was based on Cohen (2014). The optimal number of bins for the computation of the mutual information was estimated using the Freedman-Diaconis rule. On average, 8.94 bins were estimated to represent the data optimally (median 9.0, SD 2.14, range 6 — 13), thus the number of bins was set to 9 for all data sets and participants. The mutual information is reported in bits. A permutation test was performed to determine whether the two time series shared information. To this end, one of the two time series was shifted relative to the other by a random amount. This procedure was repeated 5000 times to create a null distribution, which was then used to estimate the p-values.

#### Estimation of signal and noise properties

The noise spectrum of the data was computed on the continuous data of the simultaneous data set before any other processing of the data was done. The power spectral density was estimated for a range of 1 Hz to 200 Hz in 1 Hz steps using the Fast Fourier Transform with a Hamming window. The signal-to-noise ratio (SNR) was estimated on the epoched data after line noise removal and artifact rejection. The SNR estimate was computed using the root-mean-square of a baseline (—100ms to −30 ms before flash onset) and active period (30 ms to 100 ms after flash onset) and then converted to decibels.

## 3 Results

We recorded retinal activity with 8 OPM sensors during the presentation of two sets of short flash stimuli. Figure 1 shows the recorded activity for the OPM sensors, averaged across participants. The figure shows the data from both sessions, one with an ERG electrode simultaneously recording retinal activity (Fig. 1A) and one with only the OPM sensors present (Fig. 1B). Both recordings clearly show the *a-wave* as a negative deflection around 25 ms and the *b-wave* as a positive deflection around 80 ms (also compare the ERG activity depicted in Fig. 2C). The clear shape of the a-wave and b-wave in the separate recording already argues against the possibility of the OPM sensors picking up artefactual activity from the ERG electrode, but shows that the OPMs directly record true retinal activity.

**Figure 1:**
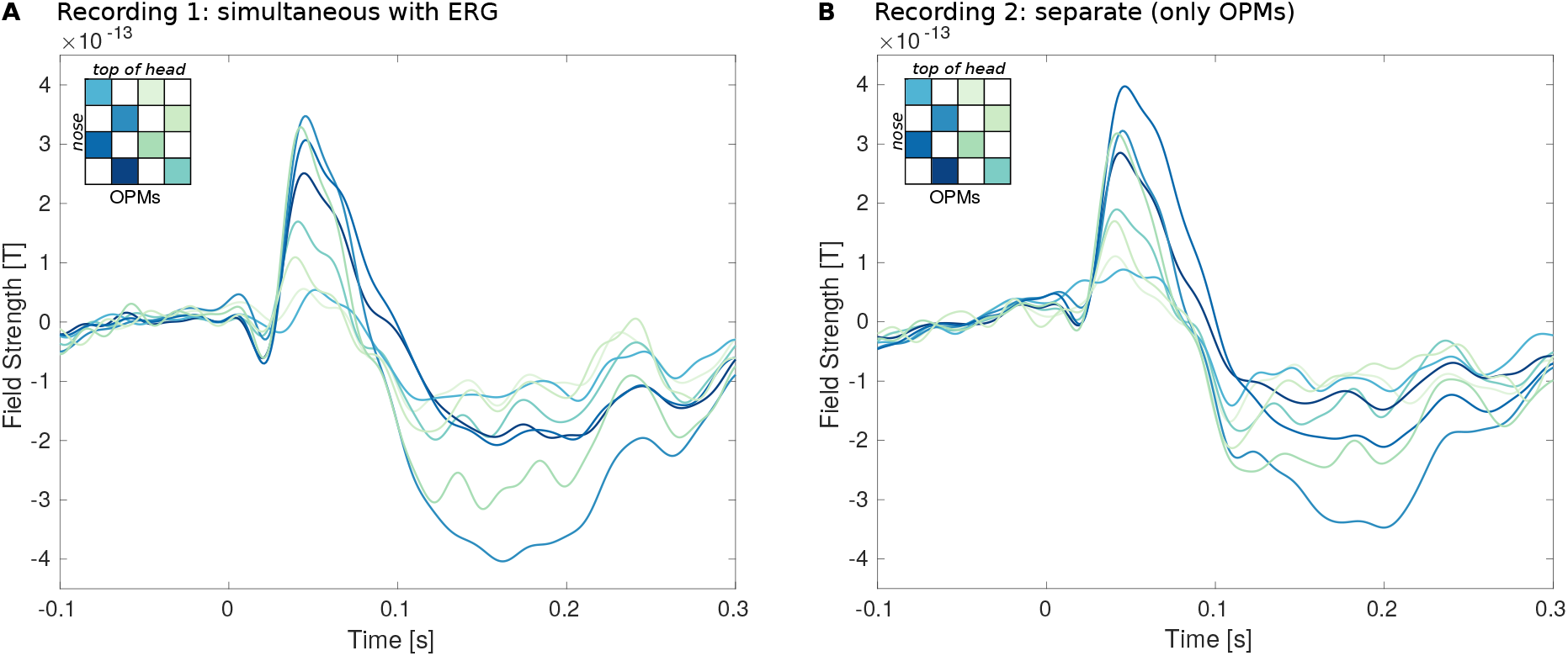
OPM recordings of retinal activity. The figure shows the two recordings of retinal activity using OPM sensors. The traces show the activity recorded by the eight OPM sensors, averaged across participants. The colors correspond to the sensors’ positions in the recording grid, depicted by the insert in the figure. **A** depicts the simultaneous recording, when the ERG electrode was also present. **B** shows the data from the second recording from the same participants after the ERG electrode was removed.

Figure 2A shows the single subject traces from the simultaneous recordings, averaged across all trials and channels. Also here, the a-wave and b-wave can be identified in all participants and gets even more pronounced when looking at the trial-averages of the best channel (defined as the channel with the highest b-wave amplitude) in Figure 2B. The ERG averages (Fig. 2) show a high resemblance to the single subject averages, but also to the data from both the simultaneous and separate recording shown in Figure 1.

**Figure 2:**
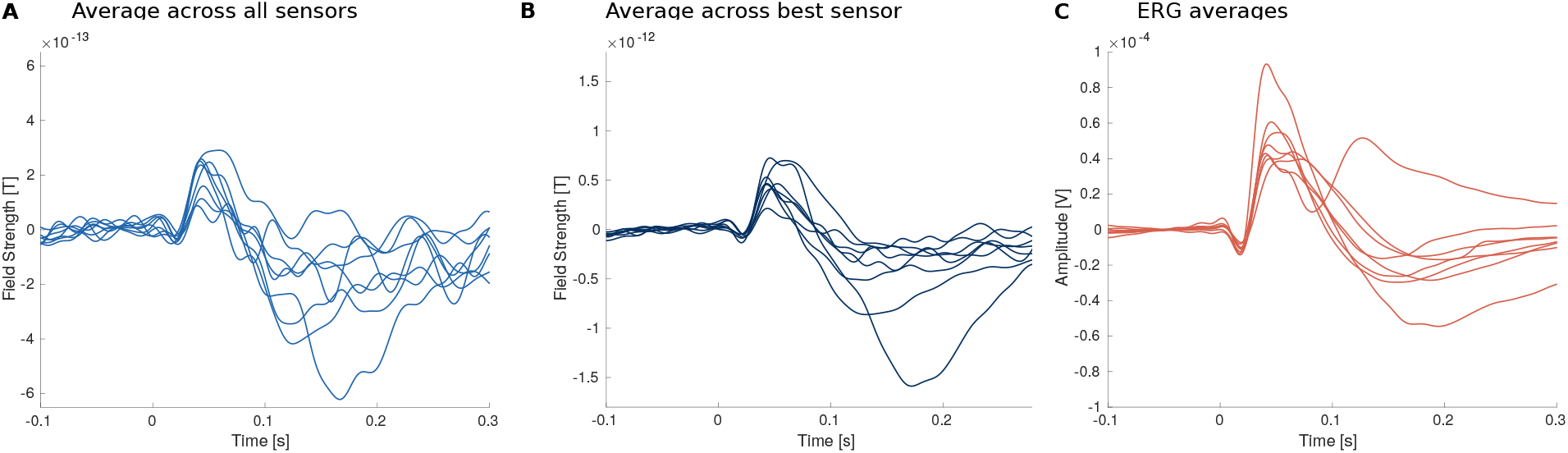
Single subject traces. Retinal activity from the simultaneous recording, each traces corresponds to one participant. **A** Per participant, the average across all trials and OPM sensors is shown. **B** Per participant, the average across all trials of the best OPM sensor is shown. The best sensor was defined individually per subject as the sensor with the highest b-wave peak amplitude. **C** Per participant, the average across trials of the ERG electrode is shown.

Figure 3A shows the ERG and MRG traces averaged across participants. For the MRG analysis, again the OPM sensor with the highest b-wave amplitude was chosen per participant and recording. The solid dark blue line corresponds to the MRG data from the simultaneous recording, while the dashed dark blue line depicts the data from the separate data set. To quantify the shared information between the ERG and MRG, mutual information was computed within subjects between the ERG electrode and the OPM sensors with the highest signal-to-noise ratio. Figure 3B shows the mutual information between the ERG and the MRG for both data sets in bits. All values are significantly higher than would be expected under the null hypothesis when no information is shared, which was determined by a permutation test (all p-values ≤ 0.0000125, Bonferroni corrected). We also tested whether the mutual information of the ERG and MRG is different between the two data sets, using a Bayesian paired samples t-test. There is no evidence for the null hypothesis that both OPM data sets exhibit the same shared information with the ERG. Surprisingly, the shared information is *higher* between the ERG and the MRG from the *separate* recording (Bayes Factor [BF] 1.711). This can be viewed as anecdotal evidence for the alternative hypothesis (Jeffreys, 1961; Lee and Wagenmaker, 2014), namely, that the shared information is not the same for the two data sets. To exclude whether this was a mere effect of the higher number of trials in the separate condition, and thus a higher SNR for the MRG evoked field, we repeated the computation of the mutual information by subsampling the MRG data for the separate condition to match the number of trials in the simultaneous condition. However, the result does not change (BF 1.755, cf. Fig. S1).

**Figure 3:**
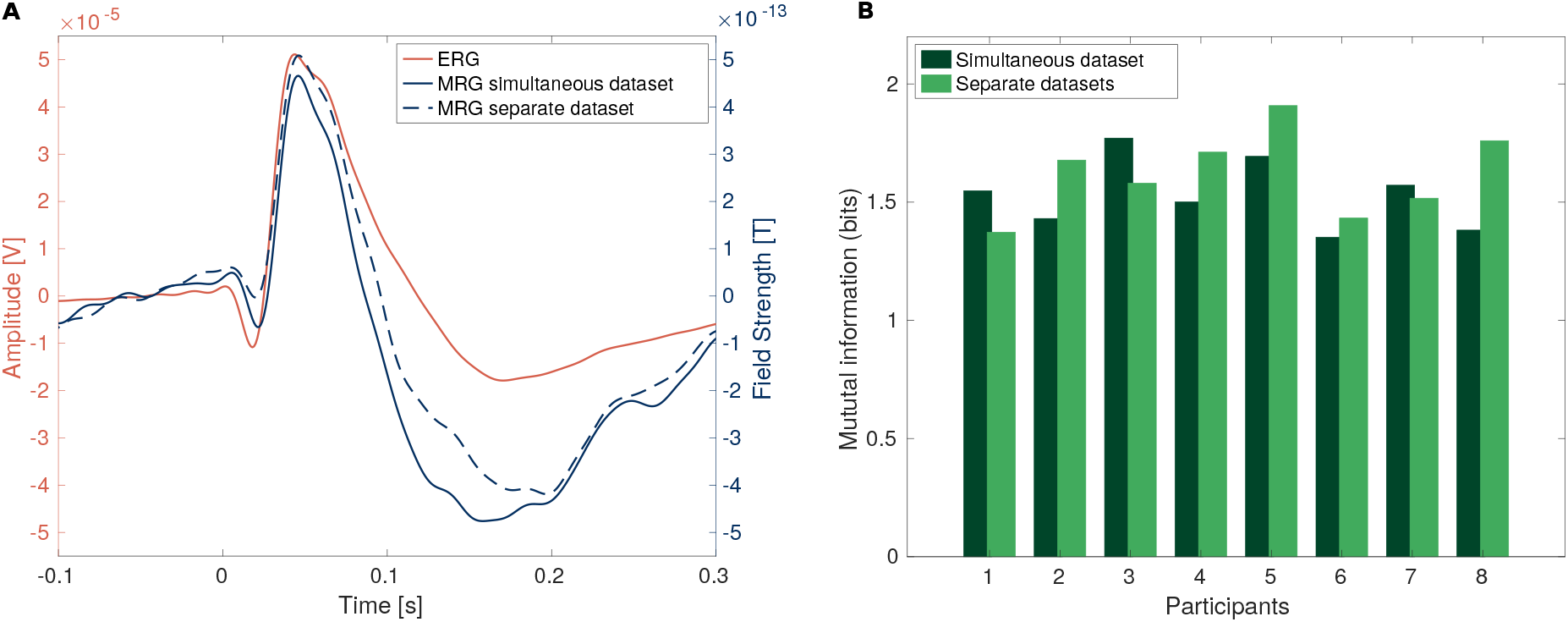
Similarity between the ERG and MRG recordings. **A** This figure shows the averages of the ERG and MRG traces across trials and participants. The orange line corresponds to the ERG electrode, the blue lines are the averages across the best OPM channel per participant (cf. Fig. 2 B) and recording (the solid line corresponds to the simultaneous recording with the ERG, and the dashed line depicts the data from the separate OPM recording). **B** The bar graph shows the mutual information between the best OPM channel and the ERG per participant. The light green bars represent the simultaneous MRG-ERG recording, and the dark green bars the separate, MRG-only recording.

We also computed the mutual information between the ERG and each OPM sensor for each participant to determine which OPM sensor positions measured retinal activity that best matched the ERG trace. This was done using the simultaneous data set. Figure 4 shows the results averaged across participants. Darker shades of green depict a higher degree of mutual information between an OPM sensor in a given position and the ERG signal. The circle size shows the precision of this estimate: the size is inversely proportional to the standard deviation of the mutual information across participants. Thus, smaller circles correspond to a less certain estimate of the mutual information. The OPM sensors in the lower left corner of the array show the highest mutual information between the MRG and the ERG, with a small standard deviation across participants. This corresponds to the sensors that were placed lateral to the nose and cheek of the participants.

**Figure 4:**
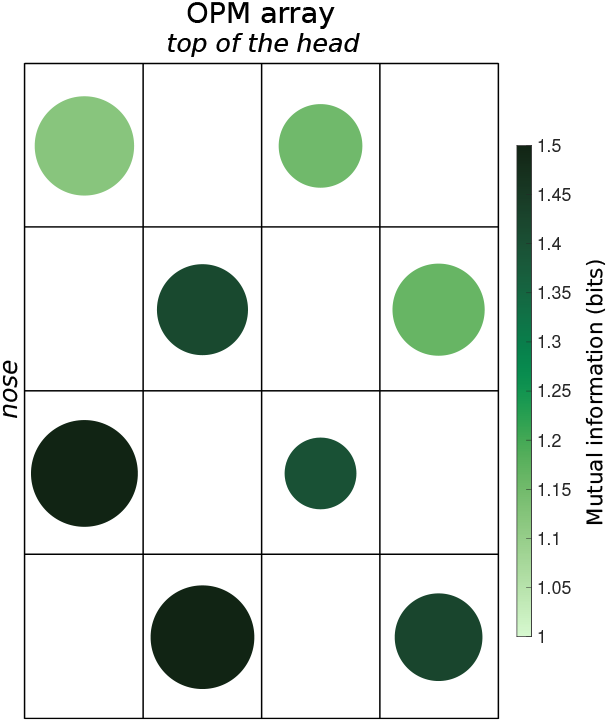
Sensitivity of the OPM array. Shown is the mutual information per channel with the ERG activity of the simultaneous recording, averaged across subjects. The color corresponds to the mutual information, the size of the circle is inversely proportional to the standard deviation across participants (such that smaller circles correspond to a less certain estimate of the mutual information).

Lastly, we looked at the signal and noise properties of both the MRG and ERG signals by computing the noise spectrum as well as SNR. Figure 5A compares the noise spectrum of the ERG (orange) and MRG (blue) across the continuous data of the simultaneous recording. Figure 5B depicts the signal to noise ratio of the single trials.

**Figure 5:**
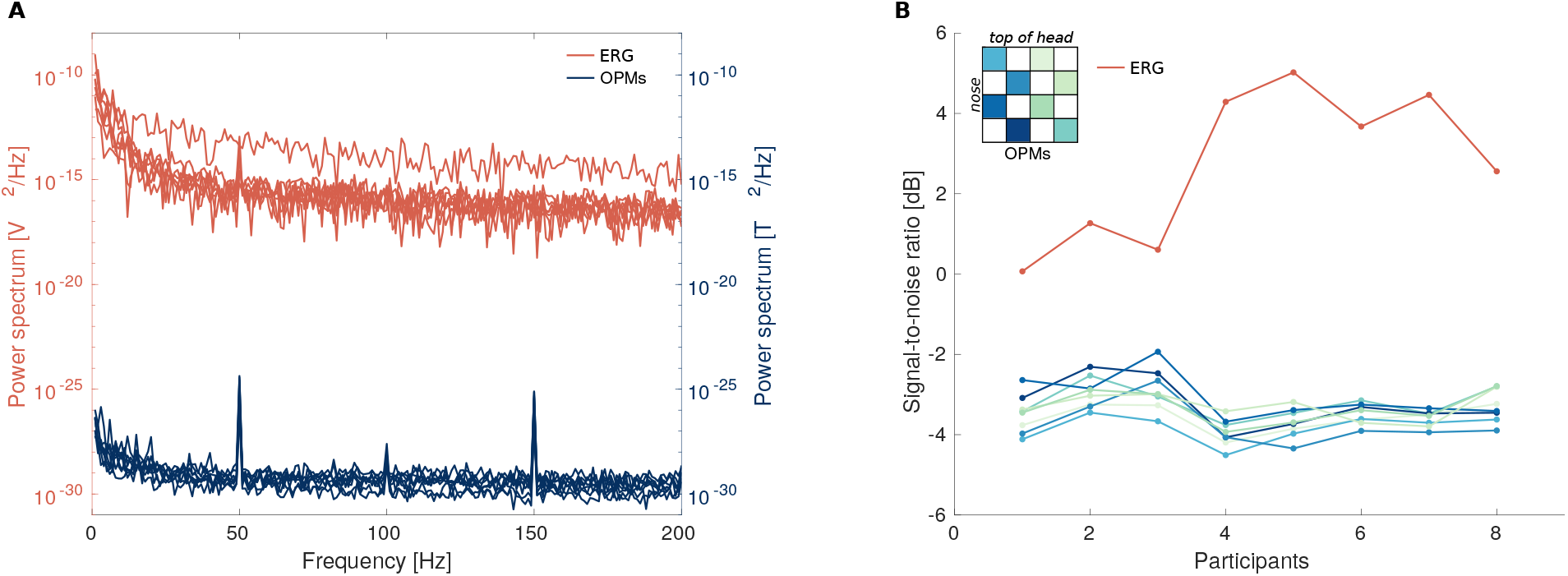
Signal and noise properties. The figure compares the signal and noise properties of the MRG and ERG recordings. **A** Shown is the noise spectrum of the ERG (orange) and the MRG (blue). The traces correspond to single subjects and the data originate from the simultaneous recording. **B** Signal-to-noise ratio of the ERG (orange) and MRG recording (blue). For the MRG recording, the colors correspond to the sensors’ positions in the grid (see insert). The data from the simultaneous recording are averaged across participants.

ERG data recorded with DTL fiber electrodes are often highly contaminated by blinks, since the electrode lies on the lower eye lid and easily gets moved during blinks or eye movements. This causes a higher number of trials having to be excluded from data analysis. This is much less the case with the MRG: since the OPM sensors are not in physical contact with the participant, the susceptibility to blink activity is comparable to that of SQUID MEG sensors. While generally of lower SNR, we found the OPM data to be less affected by artifacts: while we excluded on average 64.13 trials (SD 31.60) in the simultaneous recording based on the ERG alone (after also looking at the MRG data in a second step, on average 5.25 additional trials [SD 10.19] were excluded), we excluded only 19.63 trials (SD 19.73) from the separate OPM recording. A one-sided Bayesian paired samples t-test yields evidence for a difference between the number of trials rejected for the MRG and ERG data (BF 7.42), with more trials being rejected in the ERG data set (excluding the additionally rejected trials from the simultaneous OPM recording).

## 4 Discussion

In this paper, we recorded retinal activity with OPM sensors for the first time. We show that the recorded MRG activity exhibits both the *a-wave* and *b-wave* (Fig. 2), which are the typical retinal potentials following visual flash stimulation. We also demonstrate that the MRG activity closely matches the retinal activity as recorded with ERG fiber electrodes in the same participants (Fig. 3). Furthermore, we describe the distribution of the magnetic field of retinal activity over our recording array (Fig. 4) and defined the signal and noise characteristics of the ERG and MRG data (Fig. 5).

Several recent papers show OPM sensors successfully recording cortical and subcortical brain activity (e.g. Boto et al., 2018; Iivanainen et al., 2020; Zhang et al., 2020) and point to the potential of whole-head OPM systems (Boto et al., 2016; Hill et al., 2020). In this paper, we show that with OPMs, we are not restricted to (sub-)cortical brain activity, but can also easily record the magnetic activity of the retina. Our OPM recordings share a highly significant amount of information with the ERG recording and both the *a-wave* and the *b-wave* are clearly visible in the evoked activity. Compared to the seminal paper by Katila et al. (1981), who reported an a-wave to b-wave amplitude of approximately 0.1 pT, our measurements show an increase of roughly 6-fold (cf. Fig. 3A and Fig. 14 in Katila et al., 1981).

Surprisingly, the shared information between the MRG and ERG was slightly higher for the MRG data from the separate recording compared to the MRG from the simultaneous recording. This effect was not driven by the difference in the amount of trials that was removed due to artifacts, which was higher in the simultaneous recording. Possibly, this result can be explained by an on average slightly more favourable position of the OPM sensors for recording retinal activity in the separate recording or by a slightly poorer SNR in the simultaneous recording through artifacts induced by tiny motions of the ERG cable. Also note, that the Bayes Factor of this result is not very high, and is only classified as “anecdotal evidence” (Jeffreys, 1961; Lee and Wagenmaker, 2014). Ultimately, we can still conclude that the retinal activity in the OPM sensors is not simply induced by the ERG electrode, but genuine recorded activity.

In our experiment, the OPMs were evenly spaced out in a plastic holder that was brought lateral to the eye of the participant. The sensitivity of the grid to the retinal activity (as compared to the activity measured with the ERG electrode) was highest towards the nose and cheek of the participants. This matches the early report of MRG recordings by Katila and colleagues (1981), who also found the area below the eye to be the most sensitive to retinal activity in their recordings with a SQUID magnetometer (cf. Fig. 15 in their paper).

We placed the grid with the sensors in reference to the participants’ eyes, thus, the exact placement naturally may have varied across participants. With more exact placement, a holder for the OPMs that follows the anatomy more closely, and co-registration of the OPMs to facial landmarks, the signals should be even more comparable across participants. Then, it would even be possible to look at the topographical distribution of the signal, e.g. in combination with multifocal ERG stimulation (Derafshi et al., 2017).

While the OPM recordings have an overall lower SNR, they show less artifacts. We report a significant difference between the MRG and the ERG recordings for the number of trials that had to be rejected due to eye blinks or other artifacts. The ERG electrode has direct contact to the lower eye lid, and thus gets inevitably moved during blinks or eye movements, causing large artifacts or even signal saturation. Since the OPM array did not have any physical contact with the participant, blinks or eye movements do not cause the same disturbances in the signal. This is a clear advantage of the OPM-MRG over the fiber electrode ERG.

The SNR of the OPM-MRG recordings could be improved by further decreasing the noise in the shielded room, the background fields in our experiment were in the range of up to 15 nT. A placement of the OPM sensors around the eye that is better adjusted to the head’s shape could further increase SNR. Moreover, new developments in adaptive noise canceling (Boto et al., 2018; Iivanainen et al., 2019), one-person shielding solutions (Borna et al., 2020) or new sensor types (Zhang et al., 2020) will make it easier to record neural magnetic fields with OPM technology.

The retinal activity following flash onset also comprises a high frequency response, the oscillatory potential. Although not visible in our data set, we are optimistic about being able to record this fast oscillation with OPMs, especially in a less noisy environment. OPMs have recently been shown to capture visual gamma activity from occipital cortex up to 70 Hz (Iivanainen et al., 2019). While OPM sensors for MEG applications are limited in their bandwidth, often to 0 Hz to 200 Hz (Knappe et al., 2019), in a less noisy environment it should still be possible to record the oscillatory potential, which is centered around 120Hz (Frishman, 2013).

DTL fiber electrodes are often indicated for clinical examinations in children and sensitive adults, and have been shown to have comparable signal quality to Burian-Allen contact lens electrodes (Dawson et al., 1979; Kuze and Uji, 2000). However, even DTL electrodes require contact with the sclera, which some patients and healthy participants may still find disagreeable (Yin and Pardue, 2004; García-García et al., 2016). With further refinement of cost-effective shielding solutions (Boto et al., 2018; Iivanainen et al., 2019; Borna et al., 2020), OPMs will likely become more economically feasible in clinical settings as well. OPM sensors therefore have the potential to become a contactless alternative to ERG electrodes. The ease of obtaining multichannel coverage around the eye could also add further precision to the multifocal retinogram by enabling retinal source localization.

Furthermore, a contactless solution for recording retinal activity will benefit neuroscientific research on the human visual system. Looking ahead towards whole-head OPM systems, it would then be trivial to add some OPM channels near the eyes to capture retinal activity. This would allow the retina to be easily and comfortably examined along with the cortex in studies of visual processing.

## Acknowledgments

We would like to thank Jeramy Hughes, Orang Alem, Taylor Maydew, and Svenja Knappe from FieldLine Inc. and Vladislav Gerginov from University of Colorado for their outstanding support. Further, we would like to thank Petteri Laine from MEGIN for providing the Helmholtz coil for the MEG dewar. This research was supported by the European Union’s Horizon 2020 Marie Skłodowska Curie Individual Fellowship (grant agreement 893912) awarded to BUW. This work was supported by an ERC Starting Grant (ERC-StG-640448) to SSD. MJ was supported by a grant from the Carlsberg Foundation, grant number CF16-0004.

## Supplementary Material

**Figure S1:**
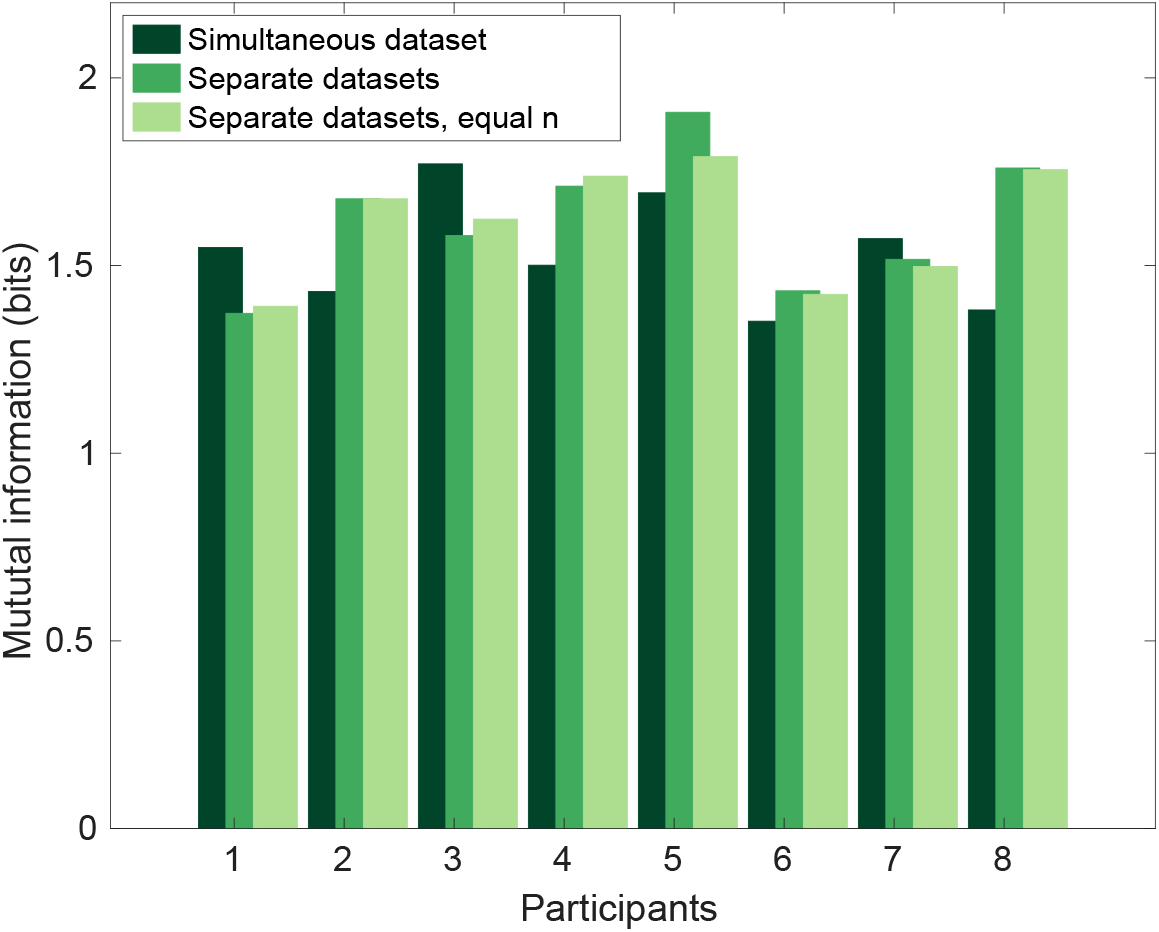
Mutual information between the ERG and MRG recordings. This figure shows the mutual information between the best OPM channel and the ERG per participant. The dark green bars represent the simultaneous MRG-ERG recording, the mid green bars the separate, MRG-only recording. The light green bars represent the MRG-only recording after randomly subsampling the data to include the same number of trials as the ERG data from the simultaneous data set.

## References

Aittoniemi K, Katila T, Kuusela ML, Varpula T (1979) In XII International Conference on Medical and Biological Engineering, Ch 94.6 (Jerusalem).

Alem O, Benison AM, Barth DS, Kitching J, Knappe S (2014) Magnetoencephalography of Epilepsy with a Microfabricated Atomic Magnetrode. Journal of Neuroscience 34:14324–14327.

Bach M, Cuno AK, Hoffmann MB (2018) Retinal conduction speed analysis reveals different origins of the P50 and N95 components of the (multifocal) pattern electroretinogram. Experimental Eye Research 169:48–53.

Borna A, Carter TR, Colombo AP, Jau YY, McKay J, Weisend M, Taulu S, Stephen JM, Schwindt PDD (2020) Non-Invasive Functional-Brain-Imaging with an OPM-based Magnetoencephalography System. PLOS ONE 15:e0227684.

Boto E, Bowtell R, Krüger P, Fromhold TM, Morris PG, Meyer SS, Barnes GR, Brookes MJ (2016) On the Potential of a New Generation of Magnetometers for MEG: A Beamformer Simulation Study. PLOS ONE 11:e0157655.

Boto E, Holmes N, Leggett J, Roberts G, Shah V, Meyer SS, Muñoz LD, Mullinger KJ, Tierney TM, Bestmann S, Barnes GR, Bowtell R, Brookes MJ (2018) Moving magnetoencephalography towards real-world applications with a wearable system. Nature 555:657–661.

Boto E, Meyer SS, Shah V, Alem O, Knappe S, Kruger P, Fromhold TM, Lim M, Glover PM, Morris PG, Bowtell R, Barnes GR, Brookes MJ (2017) A new generation of magnetoencephalography: Room temperature measurements using optically-pumped magnetometers. NeuroImage 149:404–414.

Cohen MX (2014) Analyzing Neural Time Series Data: Theory and Practice MIT press.

Dawson WW, Trick GL, Litzkow CA (1979) Improved electrode for electroretinography. Investigative Ophthalmology & Visual Science 18:988–991.

Derafshi Z, Kunzer BE, Mugler EM, Rokhmanova N, Park DW, Tajalli H, Shetty K, Ma Z, Williams JC, Hetling JR (2017) Corneal Potential Maps Measured With Multi-Electrode Electroretinography in Rat Eyes With Experimental Lesions. Investigative Ophthalmology & Visual Science 58:2863–2873.

Doty R, Kimura D (1963) Oscillatory potentials in the visual system of cats and monkeys. The Journal of Physiology 168:205.

Frishman LJ (2013) Retina chapter Electrogenesis of the Electroretinogram, pp. 177–201. Saunders.

Fröhlich F (1914) Beiträge zur allgemeinen Physiologie der Sinnesorgane. Zeitschrift für Psychologie und Physiologie der Sinnesorgane 48:28–164.

García-García Á, Muñoz-Negrete FJ, Rebolleda G (2016) Variability of the multifocal electroretinogram based on the type and position of the electrode. Documenta Ophthalmologica 133:99–108.

Heinrich SP, Bach M (2004) High-frequency oscillations in human visual cortex do not mirror retinal frequencies. Neuroscience Letters 369:55–58.

Heinrich T, Bach M (2002) Contrast adaptation in retinal and cortical evoked potentials: No adaptation to low spatial frequencies. Visual neuroscience 19:645–50.

Hill RM, Boto E, Holmes N, Hartley C, Seedat ZA, Leggett J, Roberts G, Shah V, Tierney TM, Woolrich MW, Stagg CJ, Barnes GR, Bowtell R, Slater R, Brookes MJ (2019) A tool for functional brain imaging with lifespan compliance. Nature Communications 10:4785.

Hill RM, Boto E, Rea M, Holmes N, Leggett J, Coles LA, Papastavrou M, Everton SK, Hunt BAE, Sims D, Osborne J, Shah V, Bowtell R, Brookes MJ (2020) Multi-channel whole-head OPM-MEG: Helmet design and a comparison with a conventional system. NeuroImage 219:116995.

Iivanainen J, Stenroos M, Parkkonen L (2017) Measuring MEG closer to the brain: Performance of on-scalp sensor arrays. Neuroimage 147:542–553.

Iivanainen J, Zetter R, Grön M, Hakkarainen K, Parkkonen L (2019) On-scalp MEG system utilizing an actively shielded array of optically-pumped magnetometers. NeuroImage 194:244–258.

Iivanainen J, Zetter R, Parkkonen L (2020) Potential of on-scalp MEG: Robust detection of human visual gamma-band responses. Human Brain Mapping 41:150–161.

Jeffreys H (1961) Theory of Probability Clarendon Press, 3rd edition edition.

Katila T, Maniewski R, Poutanen T, Varpula T, Karp PJ (1981) Magnetic fields produced by the human eye. Journal of Applied Physics 52:2565–2571.

Kenyon GT, Moore B, Jeffs J, Denning KS, Stephens GJ, Travis BJ, George JS, Theiler J, Marshak DW (2003) A model of high-frequency oscillatory potentials in retinal ganglion cells. Visual Neuroscience 20:465–480.

Knappe S, Sander T, Trahms L (2019) Optically Pumped Magnetometers for MEG In Supek S, Aine CJ, editors, Magnetoencephalography: From Signals to Dynamic Cortical Networks,pp. 1301–1312. Springer International Publishing, Cham.

Kuze M, Uji Y (2000) Comparison Between Dawson, Trick, and Litzkow Electrode and Contact Lens Electrodes Used in Clinical Electroretinography. Japanese Journal of Ophthalmology 44:374–380.

Lee MD, Wagenmaker EJ (2014) Bayesian Cognitive Modeling: A Practical Course Cambridge University Press.

Lopez L, Sannita WG (1997) Magnetically recorded oscillatory responses to luminance stimulation in man. Electroencephalography and Clinical Neurophysiology 104:91–95.

Marmor MF, Arden GB, Nilsson SE, Zrenner E (1989) Standard for clinical electroretinography. Archives of Ophthalmology 107:816–819.

Marmor MF, Fulton AB, Holder GE, Miyake Y, Brigell M (2009) ISCEV Standard for full-field clinical electroretinography (2008 update). Documenta Ophthalmologica 118:69–77.

McCulloch DL, Marmor MF, Brigell MG, Hamilton R, Holder GE, Tzekov R, Bach M (2015) ISCEV Standard for full-field clinical electroretinography (2015 update). Documenta Ophthalmologica 130:1–12.

Molotchnikoff S, Dubuc M, Brunette JR (1975) Simultaneous Recordings of Visual Cortex and Superior Colliculus Field Potentials in the Rabbit. Canadian Journal of Neurological Sciences/Journal Canadien des Sciences Neurologiques 2:61–66.

Munk MHJ, Neuenschwander S (2000) High-Frequency oscillations (20 to 120 Hz) and their role in visual processing. Journal of Clinical Neurophysiology 17:341–360.

Neuenschwander S, Castelo-Branco M, Baron J, Singer W (2002) Feed-forward synchronization: Propagation of temporal patterns along the retinothalamocortical pathway. Philosophical Transactions of the Royal Society B: Biological Sciences 357:1869–1876.

Oostenveld R, Fries P, Maris E, Schoffelen JM (2011) FieldTrip: Open Source Software for Advanced Analysis of MEG, EEG, and Invasive Electrophysiological Data. Intell. Neuroscience 2011:1:1–1:9.

Perlman I (2001) Webvision: The Organization of the Retina and Visual System chapter The Electroretinogram: ERG. Online Textbook of the Visual System. University of Utah.

Roberts G, Holmes N, Alexander N, Boto E, Leggett J, Hill RM, Shah V, Rea M, Vaughan R, Maguire EA, Kessler K, Beebe S, Fromhold M, Barnes GR, Bowtell R, Brookes MJ (2019) Towards OPM-MEG in a virtual reality environment. NeuroImage 199:408–417.

Shigihara Y, Hoshi H, Zeki S (2016) Early visual cortical responses produced by checkerboard pattern stimulation. NeuroImage 134:532–539.

Sieving PA, Murayama K, Naarendorp F (1994) Push–pull model of the primate photopic electroretinogram: A role for hyperpolarizing neurons in shaping the b-wave. Visual Neuroscience 11:519–532.

Tierney TM, Levy A, Barry DN, Meyer SS, Shigihara Y, Everatt M, Mellor S, Lopez JD, Bestmann S, Holmes N, Roberts G, Hill RM, Boto E, Leggett J, Shah V, Brookes MJ, Bowtell R, Maguire EA, Barnes GR (2021) Mouth magnetoencephalography: A unique perspective on the human hippocampus. NeuroImage 225:117443.

Todorov MI, Kékesi KA, Borhegyi Z, Galambos R, Juhász G, Hudetz AG (2016) Retino-cortical stimulus frequency-dependent gamma coupling: Evidence and functional implications of oscillatory potentials. Physiological Reports 4:e12986.

Vukmanic E, Godwin K, Shi P, Hughes A, DeMarco Jr. P (2014) Full-field electroretinogram response to increment and decrement stimuli. Documenta Ophthalmologica 129:85–95.

Westner BU, Dalal SS (2019) Faster than the brains speed of light: Retinocortical interactions differ in high frequency activity when processing darks and lights. bioRxiv.

Yin H, Pardue MT (2004) Performance of the DTL electrode compared to the Jet contact lens electrode in clinical testing. Documenta Ophthalmologica 108:77–86.

Zetter R, Iivanainen J, Stenroos M, Parkkonen L (2018) Requirements for Coregistration Accuracy in On-Scalp MEG. Brain Topography 31:931–948.

Zhang X, Chen Cq, Zhang Mk, Ma Cy, Zhang Y, Wang H, Guo Qq, Hu T, Liu Zb, Chang Y, Hu Kj, Yang Xd (2020) Detection and analysis of MEG signals in occipital region with double-channel OPM sensors. Journal of Neuroscience Methods 346:108948.

